# AraC mutations that suppress inactivating mutations in the C-terminal domain of the RpoA subunit of *Escherichia coli* RNA polymerase

**DOI:** 10.1101/2020.07.15.203596

**Authors:** Dominique Belin, Filo Silva

## Abstract

In *E. coli*, transcriptional activation is often mediated by the C-terminal domain of RpoA, the α subunit of RNA polymerase. Mutations that prevent activation of the arabinose P_BAD_ promoter are clustered in a small region of the α–CTD domain around K271. To determine the target(s) of RpoA in the P_BAD_ promoter, we have isolated suppressors of *rpoA* α–CTD mutations. The suppressors map to the N-terminal domain of AraC, the main transcriptional regulator of *ara* gene expression. No mutation was found in the large DNA regulatory region between *araC* and P_BAD_, suggesting that, in this system, RpoA does not activate transcription through its direct DNA binding. One class of *araC* mutations result in substitutions in the core of the N-terminal domain suggesting that they may affect its conformation. Another class of suppressors define genetically a domain that potentially interacts with the C-terminal domain of RpoA. Surprisingly, in *rpoA*^+^ strains lacking CRP, the *araC* mutations largely restore arabinose gene expression, suggesting that they somehow strengthen the AraC:α–CTD interaction. Thus, the N-terminal domain of AraC exhibits at least three activities: dimerization, arabinose binding and transcriptional activation via RpoA.

**Importance:** Gene expression is most often controlled at the level of transcription by regulators that interact with RNA polymerase. The C-terminal domain of *Escherichia coli* RpoA is attached to the core enzyme by a flexible linker and serves as a hub that interacts with many regulators and even with DNA sites to activate transcription. Mutations in a RpoA subdomain interfere with activation of the main arabinose promoter by AraC, the regulator that either activates or represses expression of the arabinose operons. We define here genetically the target of RpoA in the main arabinose promoter. Suppressors of most RpoA mutations map to the N-terminal domain of AraC that promotes its dimerization and binds to arabinose, the inducer. Thus, our results identify a third function for this AraC domain. Some suppressors define a potential binding site for RpoA, while others, at internal residues, probably affect the conformation of the AraC domain.

## Introduction

The arabinose operons were the first system in which positive control of gene expression was genetically demonstrated (1). AraC, the product of the activator gene, binds to arabinose and stimulates transcription of the catabolic genes, *araB, araA* and *araD*, as well as that of the two arabinose import operons, *araE* and *araFGH*. However, in the absence of arabinose, AraC represses transcription from the P_BAD_ promoter. Transition between the repressor and activator states of AraC involves a large conformational change, named the light switch, that displaces one subunit of the AraC dimer from the distant *O2* operator site to the promoter proximal *I2* activator site. During the switch, the N-terminal arm of AraC leaves the C-terminal DNA binding domain and covers the arabinose binding pocket of the N-terminal dimerization domain (2, 3)(Figure 1). The P_BAD_ promoter is intrinsically very weak and AraC bound to the I2 site enhances RNA polymerase binding and open complex formation (4). In addition, CRP, the cAMP binding protein, is required for full activity of the P_BAD_ promoter (5, 6). CRP activates several bacterial promoters, and several mechanisms of activation have been described (7).

**Figure 1.**
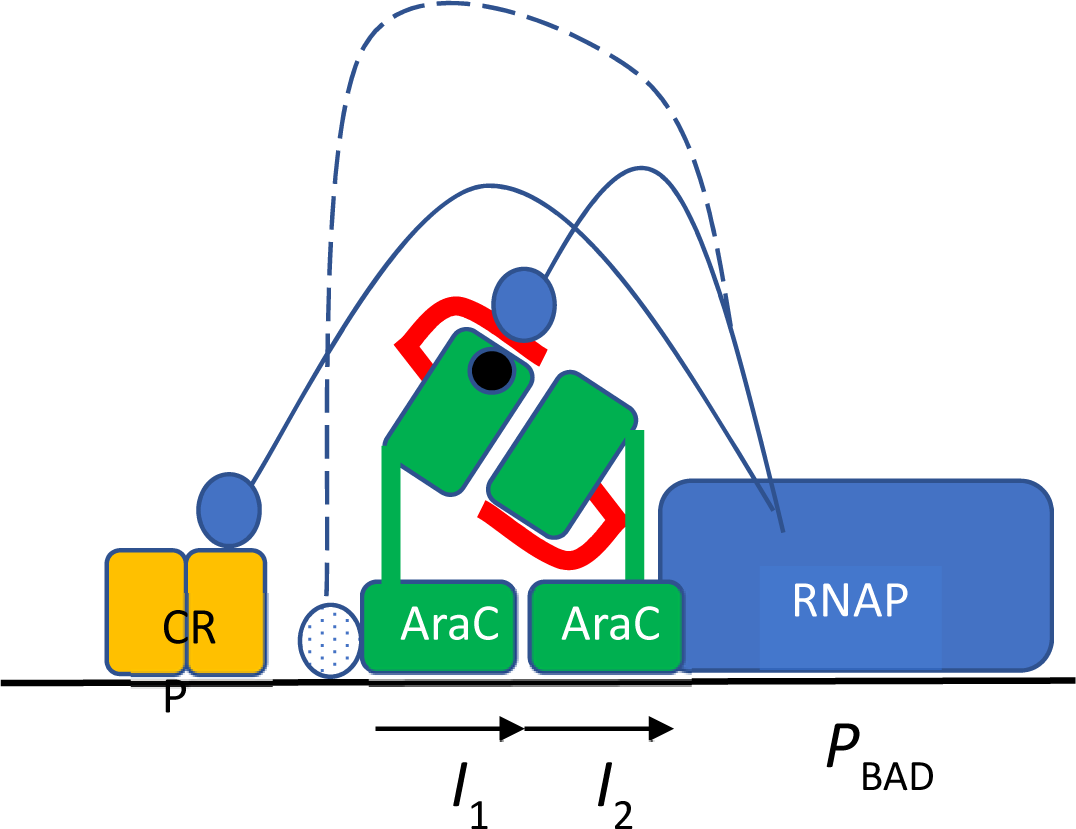
Transcription activation at the arabinose P_BAD_ promoter (adapted from (2)). RNA polymerase (blue) binding to the promoter is helped by the proximal AraC monomer bound to the *I2* site (position -35 to -51 from the start site) that overlaps the promoter -35 site. Each AraC monomer (green) consists of an N-terminal dimerization domain that binds arabinose (black circle) attached by a linker to the C-terminal DNA binding domain. In the presence of arabinose, the AraC N-terminal arm (red) covers the sugar binding pocket. The CRP dimer (yellow) binds to a distal site (position -83 to -104). One α-CTD domain of RpoA contacts CRP. The other α-CTD domain (stippled blue circle) has been proposed to contact the DNA and the DNA binding domain of the distal AraC monomer bound to *I1* (position -56 to -72). Results presented here show that the second α-CTD domain rather contacts the N-terminal arm of AraC (filled blue circle).

The α subunit of RNA polymerase, encoded by the *rpoA* gene, plays at least two roles in transcription. Its N-terminal domain (α-NTD) is involved in the assembly of the enzyme (8). In contrast, the C-terminal domain (α-CTD), which is attached by a flexible linker to the α-NTD, plays multiple roles in transcription activation (7, 9, 10). The α–CTD activates transcription by interacting with activators in different ways or by binding to UP DNA sites upstream of promoters (11, 12). Since RNA polymerase contains two α subunits, each subunit can interact independently with two activating regions (10). In the arabinose P_BAD_ promoter, one α–CTD interacts with CRP (6). The target of the second α–CTD has not yet been elucidated, but it has been proposed to bind the AraC DNA binding domain and to a DNA motif localized between the CRP and AraC binding sites (2, 13), by analogy with several CRP-dependent promoters (7). Activator mutations have been described in the phage P2 *ogr* gene using two *rpoA* alleles (14). *rpoA* α–CTD mutations that interfere with P_BAD_ activity provide a powerful genetic system to identify the interaction site(s) of α–CTD with AraC and/or a DNA element in the regulatory sequence between *araC* and *araB*. We have isolated *araC* suppressors that compensate for the detrimental effects of several mutations in the RpoA α–CTD. These mutations are scattered at several positions in the AraC N-terminal arm and dimerization domain. Some may define genetically a contact site with the RpoA α–CTD, while others rather promote or stabilize a conformation of the AraC NTD that more efficiently interacts with the *rpoA* α–CTD, thereby compensating for the deleterious effect of *rpoA* mutations.

## Materials and Methods

### Strains

DB504 is a derivative of MC4100 (15). JW5702 contains the Δ*crp*-*765*::*kan* allele (16) and was obtained from the Coli Genetic Stock Center (CGSC strain 11596). The *rpoA* mutations were introduced in DB504 by bacteriophage P1-mediated co-transduction with a *gspA*::Tn*10* marker (90-95% co-transduction) (17). The Δ*crp*-765::*kan* allele was introduced by P1-mediated transduction in strains carrying the *rpoA* alleles; the transductants were selected on streptomycin containing plates to select for recombination between the *rpsL150* marker of DB504 and *crp*, and retained the *gspA*::Tn*10* marker. Strains containing the Δ*crp*-*765*::*kan* allele were grown in LB containing 0.1% glucose (17). Information on *E. coli* genes and regulatory sequences were obtained from https://ecocyc.org/.

### Plasmids

All plasmids are derived from pBAD24 (18). pM74 expresses a maspin::PhoA chimera carrying a P32L substitution in the h-region of the signal sequence. psM74c is derived from pM74; it contains a shorter n-region and a cleaved c-region; the export of this chimeric protein is as efficient as that of wild type PhoA. pos3 contains the wild type *phoA*^+^ gene (19). A complete deletion of the *araC* coding region was performed by removing the two MfeI-digested fragments of psM74c.

### PhoA assay

PhoA activity was qualitatively detected on plates supplemented with 0.2% arabinose and 40 µg/ml XP (5-bromo-4-chloro-3 indolyl phosphate). PhoA enzymatic activity was determined by measuring the rate of *p*-nitro-phenyl-phosphate hydrolysis (20).

### Mutagenesis

Hydroxylamine mutagenesis of psM74c plasmid DNA was performed as described (19). Upon transformation in DB504, approximately 6% of the colonies carried inactivating mutations in *araC*, the regulatory region or the maspin::*phoA* gene. Plasmids allowing formation of blue colonies in Δ*crp rpoA* strains occurred at a frequency of 1-2%. More than half of the mutant plasmids exhibited a higher plasmid copy number than the original plasmid. The *araC* mutations have been independently isolated once (D7N), or three times (S14L, G22D and A152V). In a pilot experiment, an *araC* fragment was mutagenized by PCR (21) and cloned in psM74c. 10% of resulting plasmids transformed into a *rpoA*^+^ *crp*^+^ strain formed white colonies on PhoA indicator plates containing arabinose. After transformation in a *rpoA* A272E *Δcrp* strain, seven candidate suppressors were isolated and sequenced. These suppressors also suppressed the *rpoA* K271E and K271V alleles, but very poorly the N268T allele and were not further characterized.

## Results

### RpoA mutations in the α–CTD region that interfere with P_BAD_ expression

Twenty-two independent *rpoA* mutants were selected as colony-forming suppressors of the toxicity mediated by three different proteins expressed from the arabinose P_BAD_ promoter. 17 suppressors were isolated as suppressors of PAI2::PhoA that contains the signal sequence of PAI-2 fused to the mature portion of PhoA. Expression of this chimeric protein is lethal because it interferes with protein export to the periplasm (19, 21, 22). Four of the mutants suppress the toxicity of phage T4 gene *vs*.*1*, an exported lysozyme whose toxicity is weaker than that of PAI2::PhoA (23). Finally, one mutant was isolated as a suppressor of phage T4 gene *55*.*2*, a topoisomerase inhibitor that exhibit the strongest toxicity of the three proteins used (24).

The *rpoA* mutations all map to amino acids 268-272 of the RpoA C-terminal domain (α-CTD) (Figure 2A). The solution structure derived by NMR (25) is similar to that obtained from X-ray diffraction (26). The RpoA K271E substitution (*rpoA341* (27, 28)) was the most frequent one (9 isolates), but three other substitutions (V, T & I) were also identified at this position. The four K271x changes and the A272E substitutions exert similar effects, although the K271T substitution has a slightly weaker effect (Figure 2B). These mutations prevent growth on minimal media in the absence of cysteine or methionine (Cym phenotype). The L270F substitution, the weakest of the *rpoA* alleles, allows growth on minimal media and does not interfere detectably with activation of the *cysA* operon by CysB (29). The N268T substitution is the strongest *rpoA* mutation described here and the only one that suppresses *55*.*2* toxicity; it prevents growth on minimal media, even in the presence of casamino acids.

**Figure 2.**
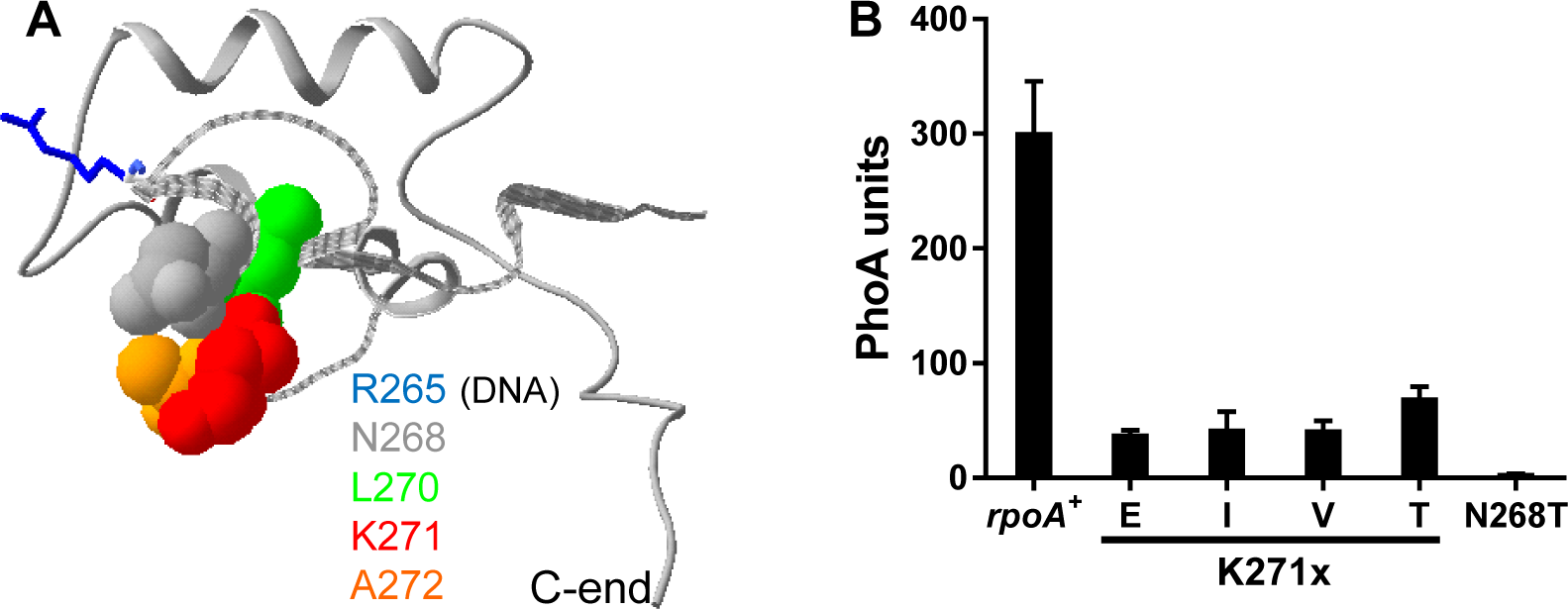
(A) The structure of the RpoA C-terminal domain as determined by NMR (PDB 1C00) (25). The image was generated with Swiss-PdbViewer v4.1.0. The domain is represented as a ribbon and the side chains of relevant residues are shown in space filling (residues 268-272) or stick (265). N268 is shown in grey, L270 in green, K271 in red and A272 in orange; R265 (blue) makes contacts with the DNA of the of the UP element λ prm and *rrnB* P1 promoters (12, 44). (B) *crp*^+^ strains containing the pM74 plasmid expressing wild type AraC and a maspin::PhoA chimera were induced for 1h with arabinose.

### Suppressors of the *rpoA* mutations

Both AraC and CRP promote transcription from the P_BAD_ promoter in the presence of arabinose (3). While AraC is absolutely required, a 20-40-fold reduced transcription can still be detected in the absence of CRP (Figure 3A). Normal P_BAD_ transcription is thought to involve interactions of the α-CTD domains of RNA polymerase with AraC and CRP (3, 6). We have measured PhoA expression from the P_BAD_ promoter in strains deleted for *crp* and carrying different alleles of *rpoA* (Figure 3B). The four substitutions of K271 strongly reduced *phoA* expression, while there was practically no expression with the N268T substitution. These findings imply that one α-CTD must interact directly with AraC, and/ or directly with DNA sites near the AraC *I1-I2* binding sites of the P_BAD_ promoter (Figure 1).

**Figure 3.**
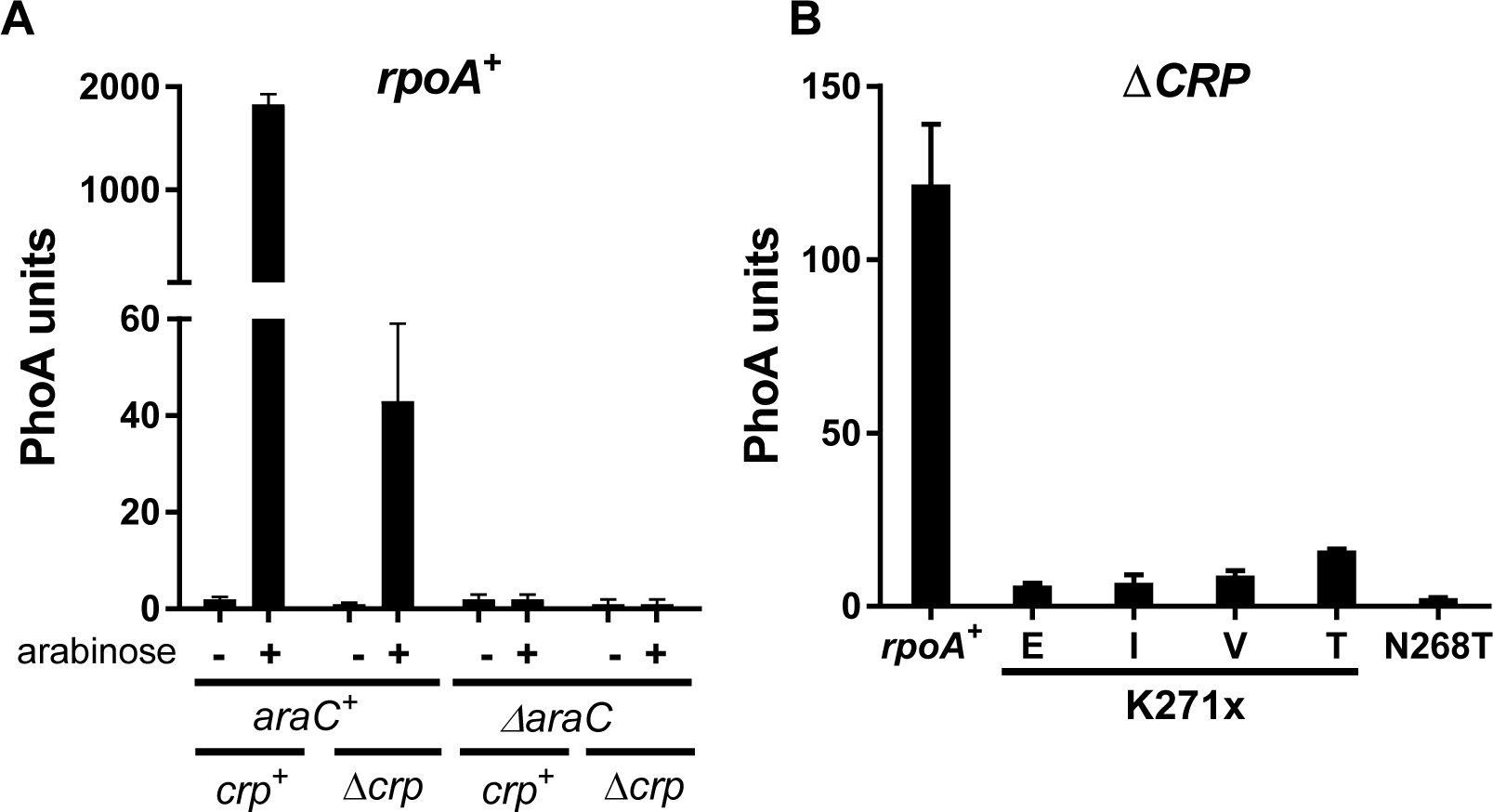
Residual activity of the P_BAD_ promoter in the absence of CRP. (A) Plasmid psM74c and its derivative with a complete deletion of *araC* were introduced in *crp*^+^ *rpoA*^+^ and Δ*crp rpoA*^+^ strains. Cultures were assayed with or without arabinose induction. (B) Pos3 expressing wild type AraC and wild type PhoA, was introduced in Δ*crp* strains carrying the indicated *rpoA* alleles. Cultures induced for 1h with arabinose

To define the interactions that are defective with the RpoA α-CTD mutants, we carried a suppressor screen. We used *Δcrp* strains that contain either the K271E or the K271V substitutions; all suppressors of either of these alleles also suppress the other substitutions of K271. The *araC* and the P_BAD_ regulatory region and promoter were present on the mutagenized pBAD plasmids. Finally, a maspin::PhoA fusion was expressed from the P_BAD_ promoter to provide a sensitive expression reporter. The maspin signal sequence of sM74c used here is highly efficient, which compensates for the reduced expression in the absence of CRP. Cells transformed with the mutagenized plasmids were plated on LB plates containing arabinose and XP, a colour indicator of PhoA activity. Plasmids isolated from blue colonies that express increased levels of PhoA were sequenced. We isolated four plasmids that contain mutations in *araC*. Mutations were also found in the P_BAD_ promoter. One, at position -35, is located between *cip*-5 site (5) and the *I*^c^ mutations (30, 31). The *cip*-5 and *I*^*c*^ promoters allow AraC independent constitutive expression and this mutation could affect RNA polymerase binding. The second one, which has a weaker stimulatory effect, is at position -23. The last one is located in the TATA box, at position - 10, and is identical to the previously isolated X^c^ mutation that creates a strong AraC-independent promoter (32, 33); a P_BAD_ promoter carrying the X^c^ mutation is not affected by the *rpoA341* K271E mutation (34). No mutation was isolated in the regulatory region between the *araC* coding sequence and the P_BAD_ promoter (Figure 1).

The *araC* suppressors of the *rpoA* α-CTD mutations localize to the NTD of AraC. Since several *araC* constitutive mutations also map to this region (35), we have measured expression from the P_BAD_ promoter in the presence and in the absence of arabinose with these AraC mutant proteins (Figure 4). In *rpoA*^+^ *Δcrp* strain, the A152V substitution confers a nearly complete constitutive phenotype, the expression measured in the absence of arabinose being 80% of that measured in the presence of arabinose. In contrast, G22D and S14L only confer a 12-27% level of expression in the absence of inducer. Constitutive expression with S14L was also observed in *crp*^+^ strains (35). Finally, the D7N substitution is entirely devoid of constitutive activity. While expression mediated by wild type AraC is strongly decreased in the absence of CRP, the *araC* mutations confer a robust expression, about 30-50% of that measured in an *araC*^+^ *rpoA*^+^ *crp*^+^ strain, which nearly compensates for the absence of CRP in *rpoA*^+^ strains (Figure 4A & C). In *rpoA* K271E strains (Figure 4B & D), the four AraC substitutions strongly promote expression in the presence of arabinose, when compared to wild type AraC. Of the four mutants tested, only the A152V one confers a partial constitutive phenotype (34% of the induced level). Thus, an arabinose constitutive phenotype is not necessary for suppression of the *rpoA* mutations.

**Figure 4.**
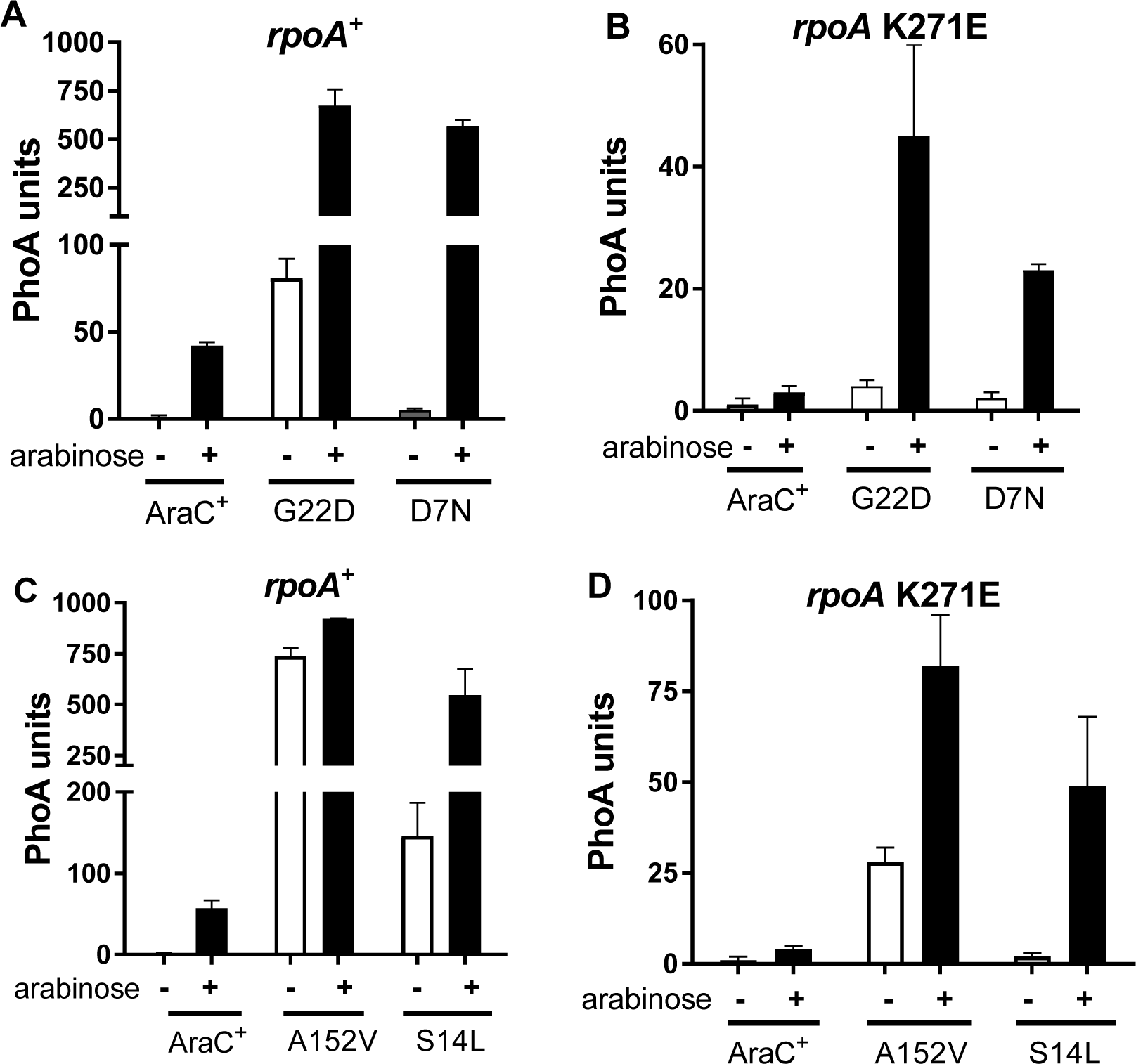
Constitutive P_BAD_ activity mediated by mutant AraC proteins in the absence of CRP. Plasmids expressing AraC^+^ or the indicated AraC suppressor proteins were introduced in Δ*crp rpoA*^+^ (A & C) or in Δ*crp rpoA341* (K271E) (B & D) strains. Cultures grown in the absence of arabinose were assayed to detect the constitutive activity of each AraC protein. Cultures induced for 1h with arabinose were used to determine the full activity of each AraC protein.

We then determined the capacity of the *araC* mutations to suppress the four different K271x substitutions, as well as N268T. The results shown in Figure 5 were obtained with AraC D7N, S14L and A152V; similar results were obtained with G22D (data not shown). In induced *rpoA*^+^ *crp*^+^ strains, the *araC* mutations had no significant effect on expression from the P_BAD_ promoter (Figure 5A, C & E). With the K271x substitutions, the increased activity was 1.5-3-fold; the highest stimulation, almost 6-fold, was observed the AraC S14L – RpoA K271E pair Figure 5C). In almost all cases, the expression observed with the *rpoA* mutants did not reach that in *rpoA*^+^ strains, indicating that the AraC suppressors do not fully compensate for the *rpoA* deficiency. The N268T substitution in *rpoA* exerts a very drastic effect on P_BAD_ expression. In this case, the *araC* mutations do stimulate expression about 3-fold (Figure 5A, C and E). In *Δcrp* strains, the stimulatory effect of the *araC* mutations was much higher (Figure 5B, D and F). In *rpoA*^+^ *Δcrp* strains, the suppressors restored expression to 30-60% of that observed in *crp*^+^ strains, nearly compensating for the absence of CRP. With *rpoA* K271x substitutions, the *araC* mutations increased expression 8-20-fold, except for K271T mutation where the increased expression was ∼30-fold. In no case was expression as high as that detected in *rpoA*^+^ strains, confirming that the *araC* mutations only partially compensate for the *rpoA* defects. The highest expression was observed with AraC A152V and RpoA K271T (Figure 5F). Finally, N268T was poorly suppressed by all the *araC* mutations.

**Figure 5.**
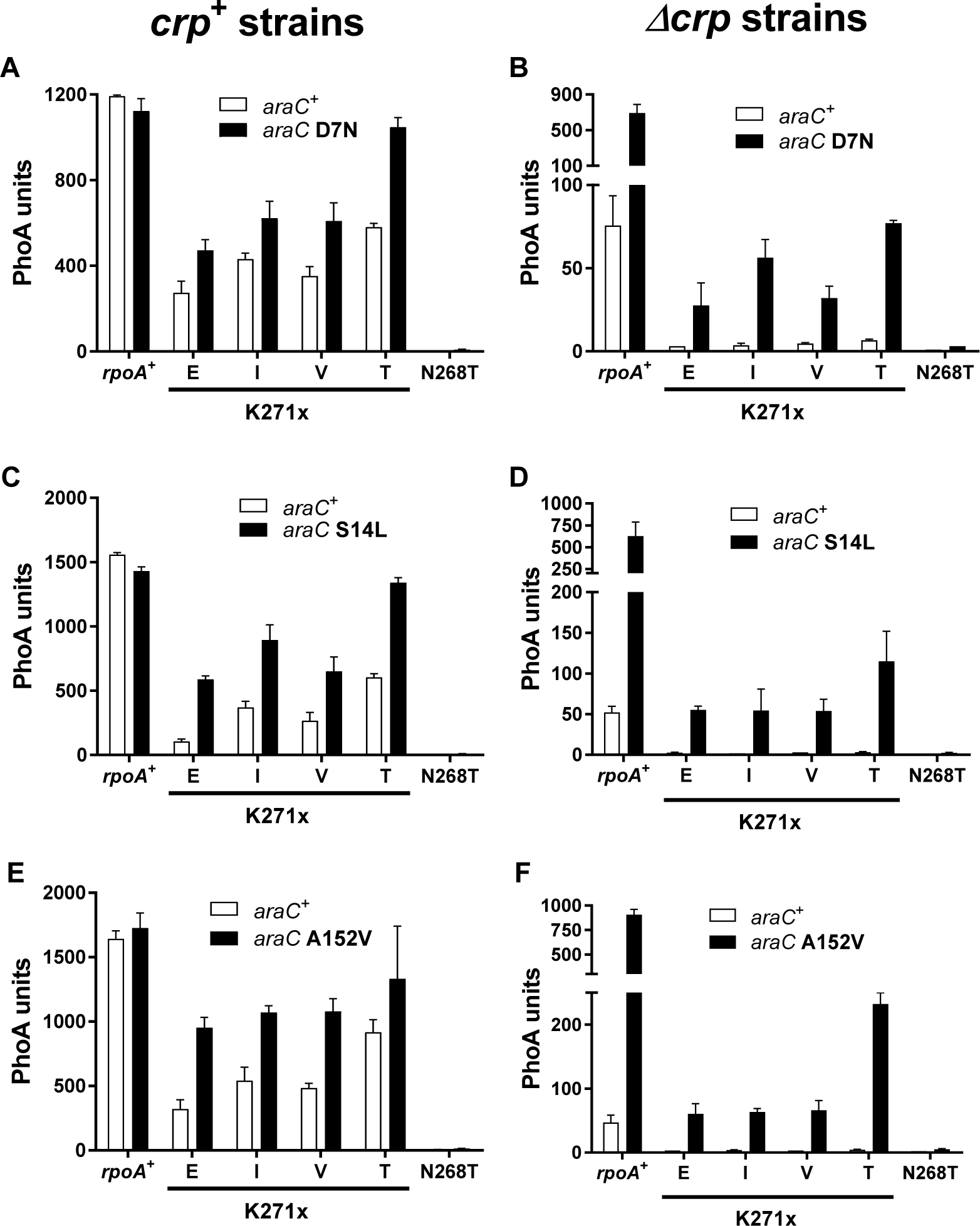
P_BAD_ activity mediated by wild type AraC^+^ and D7N (A & B), S14L (C & D) or A152V (E & F) mutant AraC proteins. Plasmids expressing AraC^+^ or the indicated AraC suppressor proteins were introduced in *crp*^+^ (A, C & E) or Δ*crp* (B, D & F) strains with the indicated *rpoA* alleles. Cultures were assayed after a 1h induction with arabinose.

To determine whether some allele specificity could be observed between the AraC suppressors and selected α–CTD mutated residues, we have compared the K271E substitution, with substitutions at the two flanking residues, L270 and A272 (Figure 6A). The four mutant AraC proteins exert similar effects in all three cases, although small quantitative differences were detected. Suppression was most efficient with the *rpoA* L270F mutant, the weakest *rpoA* substitution, and least efficient with the *rpoA* A272E mutant. Increased expression with the *araC* mutants was stronger in *Δcrp* (Figure 5A) than in *crp*^+^ (Figure 6B) strains. We did not isolate an *araC* mutation that selectively targeted a single *rpoA* allele.

**Figure 6.**
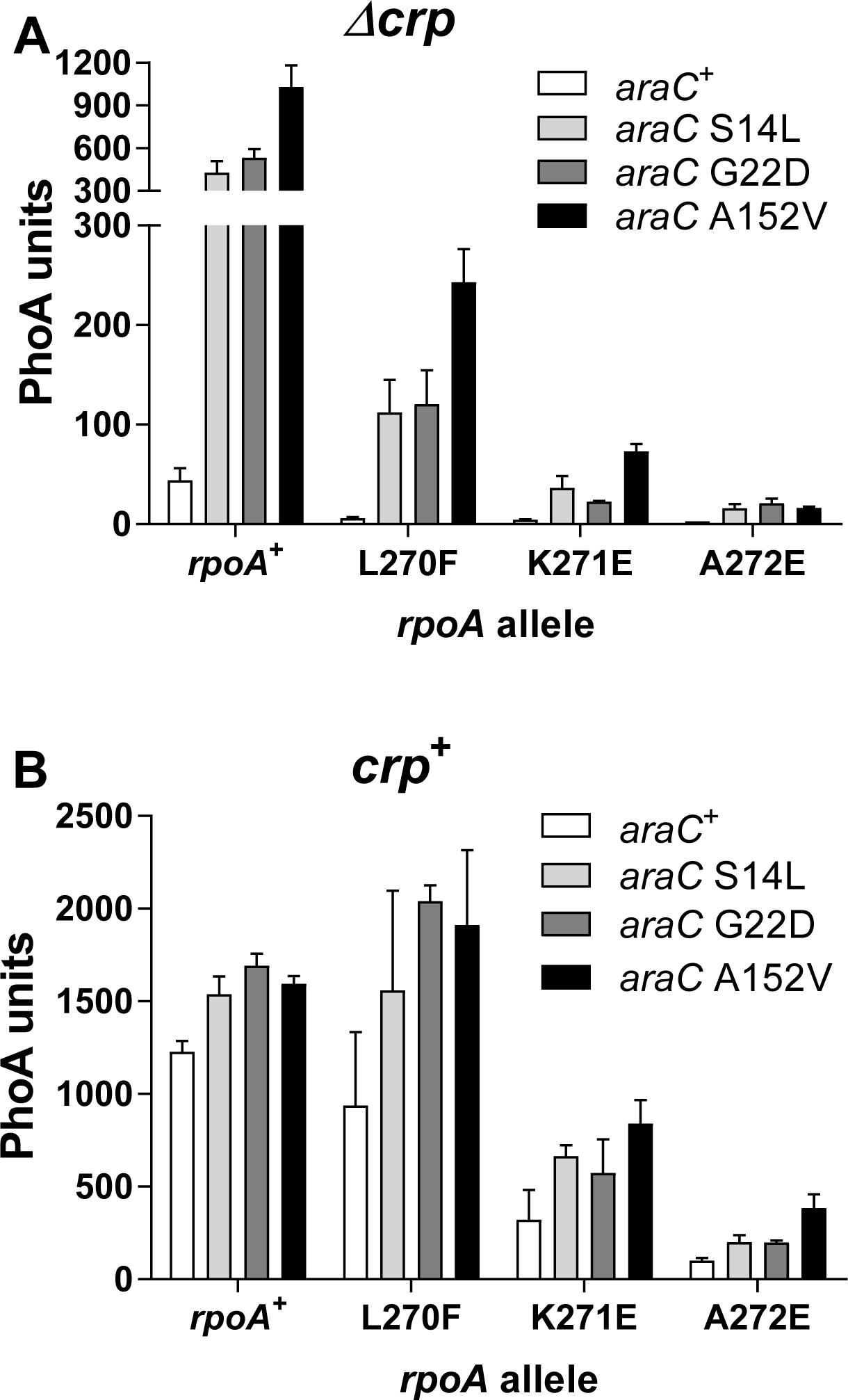
Effects of the AraC suppressors on the two residues that flank K271 of the RpoA α–CTD. Plasmids expressing AraC^+^ or the indicated AraC suppressor proteins were introduced in Δ*crp* (Figure 5A) or *crp*^+^ (Figure 5B) strains carrying the three indicated *rpoA* alleles. Cultures were assayed after a 1h induction with arabinose.

## Discussion

It was previously known that that the RpoA α–CTD domain is a pleiotropic hub required for transcription activation in multiple systems (7, 9). The residues that presumably interact with each activator are usually different. For instance, the E261K/G substitutions strongly affect a subset of CRP-dependant promoters while P322S affects OmpR (36). The S266P change allows more efficient growth on acetate medium and alters the transcription of less than 10 genes (37). The α–CTD was recently shown to play a role in genome-wide transcription regulation in *B. subtilis* (38). The K271E substitution is unusual since it also affects CysB and MelR activity in addition to AraC (34). Furthermore, we have also observed a 3-5-fold decrease of MalE synthesis in strains carrying the *rpoA* K271E substitution (data not shown); it had no effect on *malPQ* transcription, which is independent of CRP (34).

The A272T substitution has been shown to decrease *ompF* transcription mediated by OmpR (39). This mutation had no detectable effect on activation by AraC, CysB and MelR, unlike the A272E change described here that has similar effects as K271E. It is likely that the replacement of the wild type A residue by a large charged one confers a more drastic effect on α–CTD mediated transcription. With the exception of L270F, the other *rpoA* alleles described here (Figure 2A) affect MelR and CysB activation, as indicated by colony colour on MacConkey melibiose indicator plates and by their Cym phenotype (data not shown). The RpoA N268D substitution decreased *lacZ* expression about 2-fold, presumably because of a defective interaction with CRP (40). It is therefore possible that the N268T substitution described here affects activation of the P_BAD_ promoter by interfering with both AraC and CRP. This drastic mutation may interfere with the proper folding of the α–CTD or with its stability, since it affects at least one more promoter than the substitutions at the K271 and A272 sites. Indeed, all suppressors of the N268T mutation mapped to *rpoA* (data not shown) and we did not isolate an efficient suppressor in *araC*. Taken together, our results indicate that the AraC interacting region of RpoA extends at least from residues 270 to 272.

It has been previously proposed that transcription activation at P_BAD_ involves the interaction of one of the α–CTD with CRP, while the other α–CTD binds to the DNA binding domain of AraC, to a DNA sequence element between the CRP and AraC binding sites and possibly to CRP (2, 41) (Figure 1). This model is based on transcriptional activation at the *lac* promoter (7). However, the α–CTD residues involved in P_BAD_ activation are different and do not include R265 that binds DNA (11). Considering the number of *rpoA* mutations, spontaneous and UV-induced, detected with our genetic selections, it appears likely that most RpoA residues necessary for P_BAD_ activation have been identified. This provided the basis for the selection of suppressor mutations in the *araCBAD* region that could compensate for the RpoA defects.

To simplify the system, we used *Δcrp* strains so that only two of the regulatory elements were the targets of mutagenesis, the *araC* coding region and the intergenic regulatory sequence between *araC* and P_BAD_. Since expression is decreased in the absence of CRP, we used a highly sensitive reporter (Figure 3) for detection on indicator plates. Thus, suppressor mutations in genes expressed from the chromosome, such as potential compensatory mutations in *rpoA*, could not be identified. The collection of suppressors described here is limited for two reasons. Firstly, the number of characterized mutants is not large, although three of the four *araC* mutations were isolated at least twice. Secondly, most suppressors were isolated after hydroxylamine treatment of plasmid DNA, using a mutagen known to induce GC to AT transitions.

We have isolated two types of suppressors mutations. One type consists of mutations in the P_BAD_ promoter itself. They are believed to improve promoter efficiency and hence reduce the need for AraC activation. This is likely to compensate indirectly for the negative effect of the *rpoA* mutations. The other set of suppressors all carry mutation in the N-terminal arabinose binding and dimerization domain of AraC (at position 7, 14, 22 & 152). We have isolated a second set of suppressors after PCR mutagenesis of an *araC* DNA fragment. Three new substitutions were identified: N16S, I51V and I71V (Figure 7). In addition, two new changes, S14P and A152T, were found at previously characterized positions. Despite their limitations, our results strongly suggest that the N-terminal dimerization and ligand binding domain of AraC is the main target of the RpoA α–CTD during transcriptional activation of the P_BAD_ promoter. Indeed, no substitution was found in the C-terminal portion of AraC that contains the DNA binding domain. More importantly, we failed to identify any mutation in the regulatory region that would have defined a DNA site responsible for α–CTD binding (Figure 1). Furthermore, the *araC* mutations largely compensate (30-50%) for the activation defect caused by the absence of CRP in *rpoA*^+^ strains (Figures 4 & 5). Thus, they not only improve expression with the mutated α–CTD, but also strengthen the AraC interaction with wild type RpoA. If one assumes that one α–CTD binds to the NTD of the AraC monomer bound to *I1*, this may allow efficient transcription from P_BAD_. It is however possible that the second α–CTD also plays a role, for instance by interacting with the other AraC monomer bound to *I2* (Figure 1).

**Figure 7.**
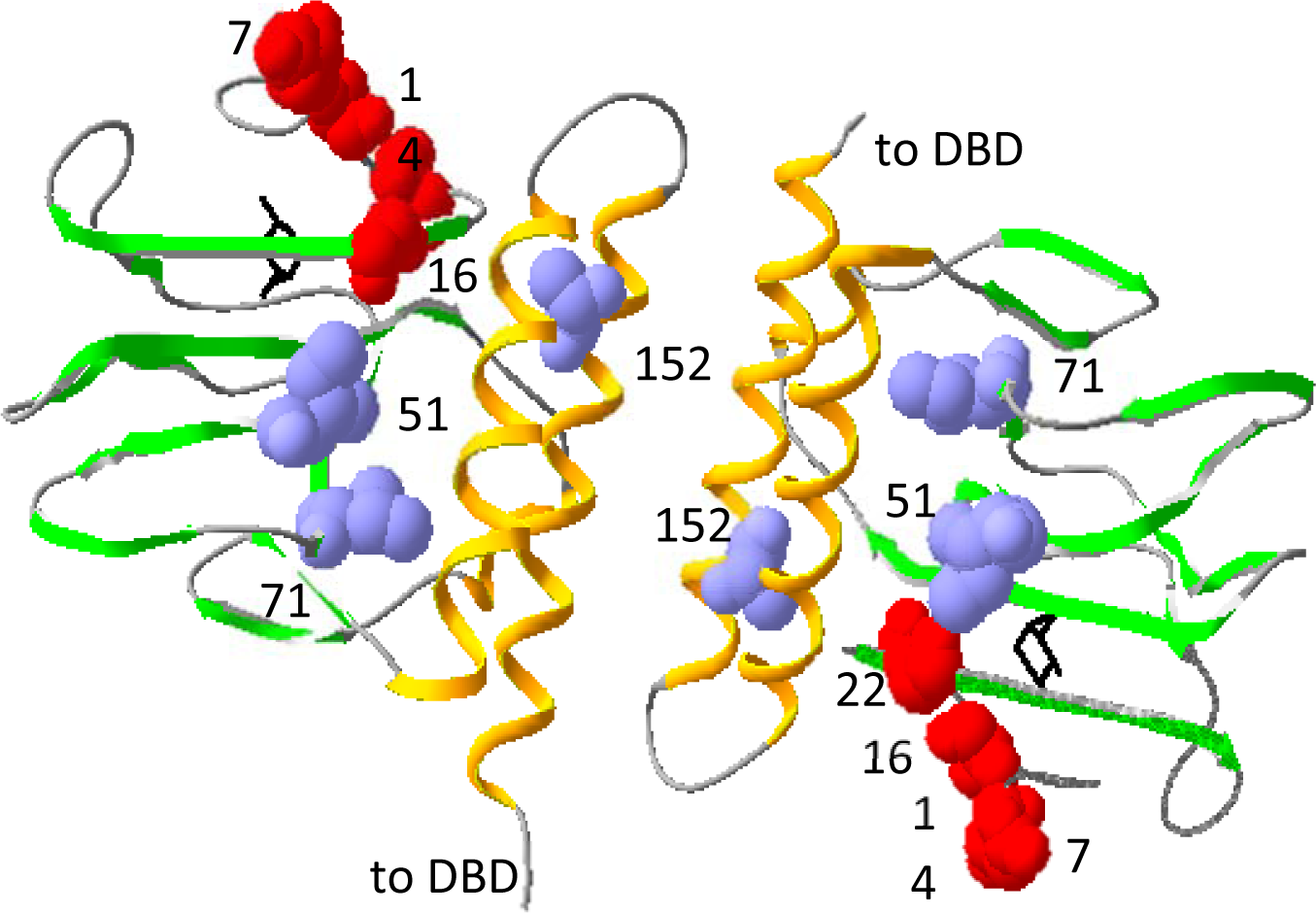
Distribution of the AraC suppressors on the structure of the AraC N-terminal domain dimer (PDB 2ARC) (42). The image was generated with Swiss-PdbViewer v4.1.0. The domain is represented as a ribbon. The two large α helices that promote dimerization are shown in orange and the β strands in green. Residues D7, S14, N16 & G22 are shown in red (N-terminal arm). I51, I71 and A152 in blue. The bound arabinose inducers are shown in black

The position of the *araC* mutations on the structure of the AraC N-terminal domain (42) are shown in Figure 7. None of the affected residues are directly involved in arabinose binding. Several arabinose constitutive mutations have been isolated in the N-terminal domain of *araC* (35, 43). The S14L, G22D and A152V mutations confer a constitutive phenotype although the first two are less active. In contrast, D7N is devoid of activity in the absence of arabinose but allowed efficient induction (Figure 4). Seven other substitutions at this site also retain an inducible phenotype (35). Thus, an arabinose constitutive phenotype is not required for *rpoA* suppression.

Several residues identified here more likely affect the structure of the AraC NTD rather than define a potential binding site for the α–CTD (Figure 7). I51 and I71 are in the core of the N-terminal domain, on the side of the β sheet that faces the two α helices that promote dimerization. A152 is in one of these helices and would also not be available for RpoA binding. This leaves D7, S14, N16 and G22 as potential elements of an α–CTD interacting region. G22 is part of the β sheet that defines the arabinose binding pocket but is close to the flexible N-terminal arm. The G22D substitution introduces a charged residue that may influence α–CTD binding either directly or indirectly. Thus, the three residues of the flexible arm, D7, S14 and N16, are potential elements of an α–CTD binding site. In agreement with this possibility, their lateral chains are closely spaced and face outside of the N-terminal domain. Furthermore, a 15 residues deletion of the arm decreased induced expression as much as several α–CTD mutations, while a 13 residues deletion has little effect (43). It is however unlikely that we have identified all the residues that could interact with the α–CTD.

We had initially hoped that our suppressor approach would have identified genetically contact sites of the RpoA α–CTD with the long regulatory region upstream of the promoter and/or with the AraC DNA binding domain, in accord with the current model of RpoA action at P_BAD_ (2). However, all the suppressors that do not change the P_BAD_ promoter map to the AraC N-terminal dimerization and arabinose binding domain. The suppressors that localize to the N-terminal arm domain suggest a new function for this highly flexible domain. In the absence of arabinose, the arm domain binds to the DNA binding domain to ensure P_BAD_ repression and DNA looping. In the presence of arabinose, the arm domain folds on top of the inducer binding pocket, and one of the arm residues, F15, contacts arabinose (35). In its third new function, the arm domain would interact with the RpoA α–CTD to promote efficient transcription at the P_BAD_ promoter. This approach may be easily extended to other activators that interact with the RpoA C-terminal domain, a hub for transcriptional activation.

## Acknowledgements

We thank Sandrine Bost, Corinne Chauffat, Giuseppe Plaia and Nessim Kaufman for the isolation of *rpoA* mutants, and Sylvie Trichard and Halil Bagci for performing some of the PhoA assays. We thank Costa Georgopoulos, Gael Panis, David Shore & Benjamin Weiss for a critical and constructive reading of the manuscript.

The work was supported by the Canton de Genève and by the CPG foundation.

